# Stimulus effects dwarf task effects in human visual cortex

**DOI:** 10.1101/2025.06.18.660183

**Authors:** Marieke Mur, Daniel J. Mitchell, Stephan Brüggemann, John Duncan

## Abstract

Task context affects stimulus representations in human visual cortex, suggesting that visual representations are flexible. However, this interpretation is at odds with a major computational goal of the human visual system: creating a perceptually stable representation of the external visual environment. How does the visual system balance stability and flexibility? Here, human participants (71 percent females) categorized object images and written words according to different task rules, while brain responses were measured with fMRI. Using an ANOVA-based modeling strategy, we precisely quantified the relative contributions of stimulus, task, and their interaction in explaining representational variance across the cortical hierarchy. Our results show that stimulus effects account for the overwhelming majority of explainable representational variance across the ventral visual system: > 95 percent in V1 and V2, and > 90 percent in higher-level visual cortex. In prefrontal cortex, the relative contributions reverse: task effects dominate stimulus effects, accounting for 80 percent of explainable representational variance. In parietal cortex, contributions of stimulus and task are approximately equal. Our findings suggest that population coding in sensory cortex is optimized for representational stability to allow a consistent interpretation of the external environment. Population coding in parietal and frontal multiple-demand cortex, by contrast, is optimized for representational flexibility to accommodate changing behavioral goals and support flexible cognition and action.

**Significance statement:** Stimulus representations in human visual cortex are affected by behavioral goals and are therefore thought to be flexible. However, this view is inconsistent with a major computational goal of the human visual system: creating a perceptually stable representation of the external environment.

Here, we show that modulatory effects of behavioral goals on stimulus representations in visual cortex are surprisingly small. In contrast, behavioral goals strongly affect representations in parietal and frontal multiple-demand cortex. Our findings suggest that population coding in sensory cortex is optimized for stable perception, while population coding in parietal and frontal multiple-demand cortex is optimized for flexible cognition.

## Introduction

It is widely thought that visual cortex supports flexible representations of stimuli. Task context affects object responses in human ventral visual cortex (O’Craven et al., 1999; Peelen et al., 2009; Stokes et al., 2009; Peelen and Kastner, 2011; Çukur et al., 2013; Harel et al., 2014; Kay and Yeatman, 2017; Nastase et al., 2017). For instance, preparatory attention for objects induces category-specific responses in high-level visual cortex, prior to the onset of the stimulus (Peelen and Kastner, 2011). Furthermore, cognitive operations required by different tasks influence category-specific responses across visual cortex, even when stimuli are held constant across tasks (Kay and Yeatman, 2017; Nastase et al., 2017). For example, attending to which animal is present in a visual scene versus what the animal is doing, enhances decodability of animal versus action information from responses in high-level visual cortex (Nastase et al., 2017). These findings are consistent with the idea that behavioral goals shape brain representations in sensory regions, enhancing objects or object properties that are behaviorally relevant.

Flexible representations are desirable because flexibility supports adaptive behavior. However, representations should also support stable perception. In fact, one of the major challenges faced by the visual system is to compute object representations that are invariant to identity-preserving image transformations (DiCarlo and Cox, 2007). Perhaps task context does not change the nature of object representations in visual cortex as dramatically as commonly assumed. Certainly, attention, driven by current behavioral goals, can selectively affect the strength of visual representations (Desimone and Duncan, 1995; Maunsell and Treue, 2006; Peelen et al., 2009). However, other brain systems, including frontoparietal multiple-demand (MD) cortex (Duncan, 2010), might introduce further and more profound representational flexibility by adaptively integrating stable perceptual input with changing behavioral goals (Roy et al., 2010; Mante et al., 2013). In line with this idea, several recent human studies have reported subtle or undetectable task effects in ventral visual regions and stronger task effects in frontoparietal regions (Bracci et al., 2017; Bugatus et al., 2017; Jackson et al., 2017; Vaziri-Pashkam and Xu, 2017).

Together, these findings raise a core question: What balance does the visual system strike between flexibility and stability? In other words, how strongly does task affect visual representations? And, how does the strength of task effects in visual regions compare to that in MD regions? Here, we address these questions by measuring functional magnetic resonance imaging (fMRI) activity in visual and MD regions while subjects perform different tasks on the same set of visual objects. We characterize object representations in visual and MD cortex across tasks using pattern-information analysis, and precisely quantify the relative contribution of stimulus, task, and their interaction in explaining the representations. We show that stimulus effects dominate task effects throughout the visual ventral stream, accounting for > 90 percent of explainable representational variance. In contrast, in the inferior frontal junction (IFJ), a core component of MD cortex, task effects dominate stimulus effects. These findings suggest that object representations in human visual cortex are surprisingly stable across task contexts.

## Materials and methods

### Subjects

Fourteen right-handed healthy native English speakers from the Cognition and Brain Sciences Unit volunteer panel participated in the fMRI experiment (age range = 21-39 years; 10 females). Subjects had normal or corrected-to-normal vision and no history of neurological disorders. Before scanning, subjects received information about the procedure of the experiment and gave written informed consent for participating. The experiment was approved by the Cambridge Psychology Research Ethics Committee, Cambridge, UK.

### Experimental design

#### Stimuli

Stimuli consisted of 16 colored images of isolated manmade objects, and 16 written words describing the objects (Figure 1, top panel). Images were sourced from the internet, masked, and resized to the same resolution (300 x 300 pixels; the largest object dimension (width or height) was scaled to 300 pixels). Words were created in CorelDraw, using a range of fonts in black print, and saved at the same resolution as the images. All words spanned the full width of the stimulus. Stimuli were displayed at the center of fixation, on a uniform grey background, at a width of 9° of visual angle.

**Figure 1.**
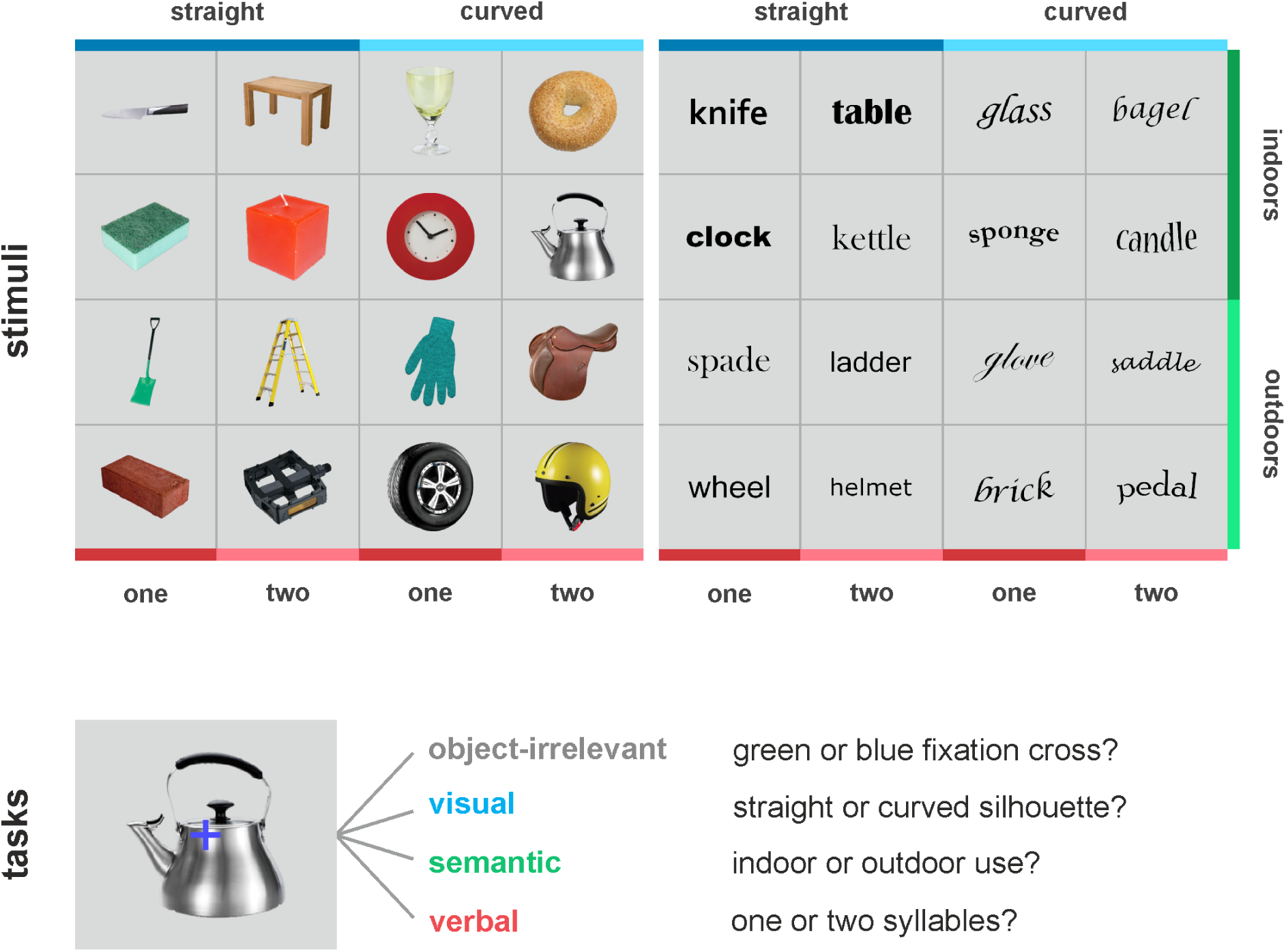
Experimental design. Stimuli consisted of 16 images of manmade objects and 16 nouns describing the manmade objects. Subjects performed four two-way categorization tasks on the stimuli: a visual task, a semantic task, a verbal task, and a task at fixation unrelated to the stimuli (“object-irrelevant”).

#### Tasks

Subjects performed four two-way categorization tasks on the stimuli (Figure 1, bottom panel). Three of the four tasks focused on object information. The first of these tasks was a visual task. For images, subjects had to indicate whether the silhouette of the image was predominantly straight or curved. For words, subjects had to indicate whether the word was written along a straight or curved line.

The second task was a semantic task. Subjects had to indicate whether the object, either shown as an image or as a word, was typically used indoors or outdoors. The third task was a verbal task.

Subjects had to indicate whether the noun describing the object consisted of one or two syllables. For images, this task required object naming; for words, this task simply required reading. The top panel of Figure 1 lists the task-relevant object information for each stimulus. For the fourth task, subjects were asked to focus on the fixation cross instead of on the objects. The color of the fixation cross randomly changed from white to either light blue or light green when a stimulus appeared.

Subjects had to indicate whether the color changed to blue or green. We refer to this task as “object-irrelevant”. The combination of 32 stimuli and four tasks yields a total of 128 conditions.

#### Design

Stimuli were presented using an event-related design, but were blocked by task. Each task block lasted approximately two minutes and started with an instruction cue, which indicated the task and response mapping for the current block. For example, for the visual task, the instruction cue showed the words “straight” and “curved”, one on either side of the fixation cross. The location of the words indicated which response button to press for each category, i.e. left index finger for the category shown on the left, and right index finger for the category shown on the right. The instruction cue was presented for a duration of 1500 ms and was followed by 4500 ms of baseline, to allow subjects to prepare for the task. Baseline consisted of a uniform gray background. Subsequently, each of the 32 stimuli appeared once in random order. Stimuli were presented for a duration of 500 ms and were followed by 2500 ms of baseline. Each task block further contained eight randomly interspersed baseline trials, which were 3000 ms in duration. Each run consisted of eight task blocks: each of the four tasks appeared once in random order, followed by 18 s of baseline providing a short rest period for the subject, after which each of the tasks appeared again in random order. Within each run half, task blocks were separated by 6 s of baseline. For each task, the left-right response mapping was randomly chosen for the first run half, and switched for the second run half. Each run started and ended with 15 s of baseline. One run lasted approximately 18 minutes in total. Subjects performed three runs per measurement session, for a total of two sessions. This yielded 12 estimates for each of the 128 conditions. Subjects were familiarized with the stimuli and tasks before scanning, and performed one training run before scanning. Subjects were instructed to be both accurate and fast.

### Functional magnetic resonance imaging

Blood-oxygen-level-dependent fMRI measurements were performed at standard resolution (voxel volume: 3 x 3 x 3 mm), using a 3T Siemens TIM Trio MRI scanner and a 32-channel head coil. We acquired 37 oblique slices (0.6 mm gap), covering the whole brain, using single-shot descending gradient echo planar imaging (EPI). Imaging parameters were as follows: EPI matrix size: 64 x 64, TR: 2000 ms, TE: 30 ms, 546 volumes per run. In addition, we acquired high-resolution T1-weighted whole-brain anatomical scans with a parallel imaging MPRAGE sequence (GRAPPA, acceleration factor 2). Imaging parameters were as follows: matrix size: 256 x 240, voxel volume: 1 x 1 x 1 mm, 192 slices, TR: 2250 ms, TE: 3 ms. We also recorded behavioral performance, consisting of reaction time and accuracy, during scanning.

### Statistical analysis

Data were analyzed using Statistical Parametric Mapping 8 (SPM8; www.fil.ion.ucl.ac.uk/spm/) and Matlab (MathWorks Inc.).

#### fMRI data preprocessing

Functional volumes from both measurement sessions were aligned to the first volume of the first session, and corrected for slice timing differences, using the middle slice as reference. The functional volumes were not spatially smoothed and remained in native space. Anatomical volumes were coregistered to the realigned mean functional volume of the first session, segmented, and normalized to MNI space. In addition, the coregistered anatomical volumes were subjected to cortical surface reconstruction in FreeSurfer (http://surfer.nmr.mgh.harvard.edu/). The normalized anatomical volumes and reconstructed cortical surfaces were used for the definition of regions of interest. Runs with excessive head motion (translation or rotation exceeding the voxel size) were excluded from analysis, leaving 87% of the data for analysis. The analyzed data consisted of all six runs for seven out of 14 subjects, five runs for five out of 14 subjects, and three runs for two out of 14 subjects.

#### Definition of regions of interest

We defined regions of interest (ROIs) in visual and multiple-demand cortex. ROIs were defined in individual subjects by selecting voxels most responsive during the experiment within predefined anatomical or functional masks.

*ROI masks.* We created anatomical masks for V1, V2, V3v, and V4 using the SPM anatomy toolbox (Eickhoff et al., 2005). The toolbox contains probabilistic maps for a range of areas, defined using cytoarchitectonic analysis of 10 human postmortem brains. Anatomical masks were created from the maximum probability map, which assigns each voxel to the area with the highest probability at that voxel. A mask for a specific area, e.g. left V1, includes the voxels assigned to that specific area and excludes all other voxels. For IT, we used an anatomical mask from Charest et al. (2014), which was manually drawn on a subject-average cortical surface. For parietal and frontal cortex, we used functional masks from Fedorenko et al. (2013). Fedorenko et al. (2013) report regions that are consistently activated across a range of cognitive demands in a group of 40 human subjects. From this set of multiple-demand regions (Duncan, 2010), we selected the intraparietal sulcus (IPS) and the inferior frontal junction (IFJ), given their presumed homology to regions in the macaque brain that represent task-dependent category information about visual stimuli, i.e. the lateral intraparietal area (LIP) and lateral prefronal cortex (lPFC), respectively (Petrides and Pandya, 1999; Brass et al., 2005; Freedman and Assad, 2006; Roy et al., 2010; Mitchell et al., 2016; Orban, 2016). Masks were then projected to native subject space. For masks in MNI space (V1, V2, V3v, V4, IPS, IFJ), we performed this step using the inverse normalization parameters obtained during segmentation and normalization in SPM8. For the IT mask, we resampled the subject-average cortical surface mask onto the native cortical surfaces and transformed the native surface masks back to volumetric space using FreeSurfer and custom matlab code freely available from https://github.com/kendrickkay.

Overlap between the V1-V4 masks and the IT mask was removed from the IT mask, and all masks were restricted to grey-matter voxels.

*ROI voxel selection.* ROIs were defined in each hemisphere by selecting, from the masks, the 71 voxels most responsive to the stimuli in the four tasks. We prevented biased results by selecting voxels based on the training data used in the main analyses (see *Construction of the representational dissimilarity matrix*). Because we used an exhaustive leave-one-run-out cross-validation procedure for the main analyses, each ROI was defined as many times as there were cross-validation folds. For each fold, we concatenated the training data, consisting of all runs except the test run, along the temporal dimension. We fit a univariate linear model to obtain a response-amplitude estimate (beta) for each of the 128 conditions at each voxel. The model included a hemodynamic response predictor for each of the 128 conditions and for each of the eight instruction cues. The predictor time courses were computed by convolving events with the SPM canonical hemodynamic response function. Event durations were set to the stimulus duration. The model further included, for each training run, six head-motion predictors, a linear-trend predictor, a 10-predictor Fourier basis for non-linear trends (sines and cosines of up to five cycles per run), and a confound-mean predictor.

For each condition, we converted the resulting beta map into a t map. Responsiveness was assessed using the t map for the average response to the 128 conditions. Of all voxels across folds that were selected for their responsiveness, 70-80 percent were selected in each fold, 15-25 percent were selected in a subset of folds, and 5 percent were selected in only one fold. These are subject-average percentages and the reported range reflects variability between ROIs.

#### Construction of the representational dissimilarity matrix

To characterize the information about the stimuli and tasks represented by a particular ROI, we computed a representational dissimilarity matrix (RDM, Kriegeskorte et al., 2008), consisting of response-pattern dissimilarities between all condition pairs. We used the linear discriminant t value (LDt, Nili et al., 2014) to estimate response-pattern dissimilarity. LDts were computed using a leave-one-run-out cross-validation procedure. For each fold, we used the training data, consisting of all runs except the test run, to determine Fisher linear discriminants for all condition pairs, and then projected the test data onto these discriminants. The LDt is the normalised contrast between two test response patterns after projection onto their discriminant. The LDt can be interpreted as a normalized cross-validated Mahalanobis distance. It provides an unbiased estimate of the discriminability of two response patterns elicited by different conditions. The LDt is sensitive to pattern scaling (Walther et al., 2016), which includes spatial-mean activation differences between conditions. LDts were averaged across folds, yielding one RDM for each ROI in each subject. RDMs are mirror-symmetric about the main diagonal. We did not compute LDts for the main diagonal because the LDt is an unbiased dissimilarity estimate and therefore the expected value for these entries is zero.

#### Descriptive visualizations

To get an impression of the stimulus representation across task contexts, we visualized the data in multipe ways. Figure 2A displays the RDMs. For each ROI, RDMs were averaged across hemispheres and subjects, and transformed into percentiles to enable easier comparison of the representational structure between ROIs. Figure 2B displays the multidimensional scaling (MDS) arrangements (Torgerson, 1958; Shepard, 1980) associated with the RDMs. The MDS arrangements (criterion: metric stress) display the multidimensional representation in two dimensions with minimal distortion; the further apart the stimuli, the more dissimilar their response patterns. MDS was performed on the non-transformed LDts; negative LDts were set to zero before applying MDS. Axes of the MDS arrangements were scaled to enable easier comparison between ROIs. For each ROI, the MDS arrangement is shown three times: once with the actual stimuli, once with the stimuli intensity-coded for object category, and once with the stimuli color-coded for task context.

**Figure 2.**
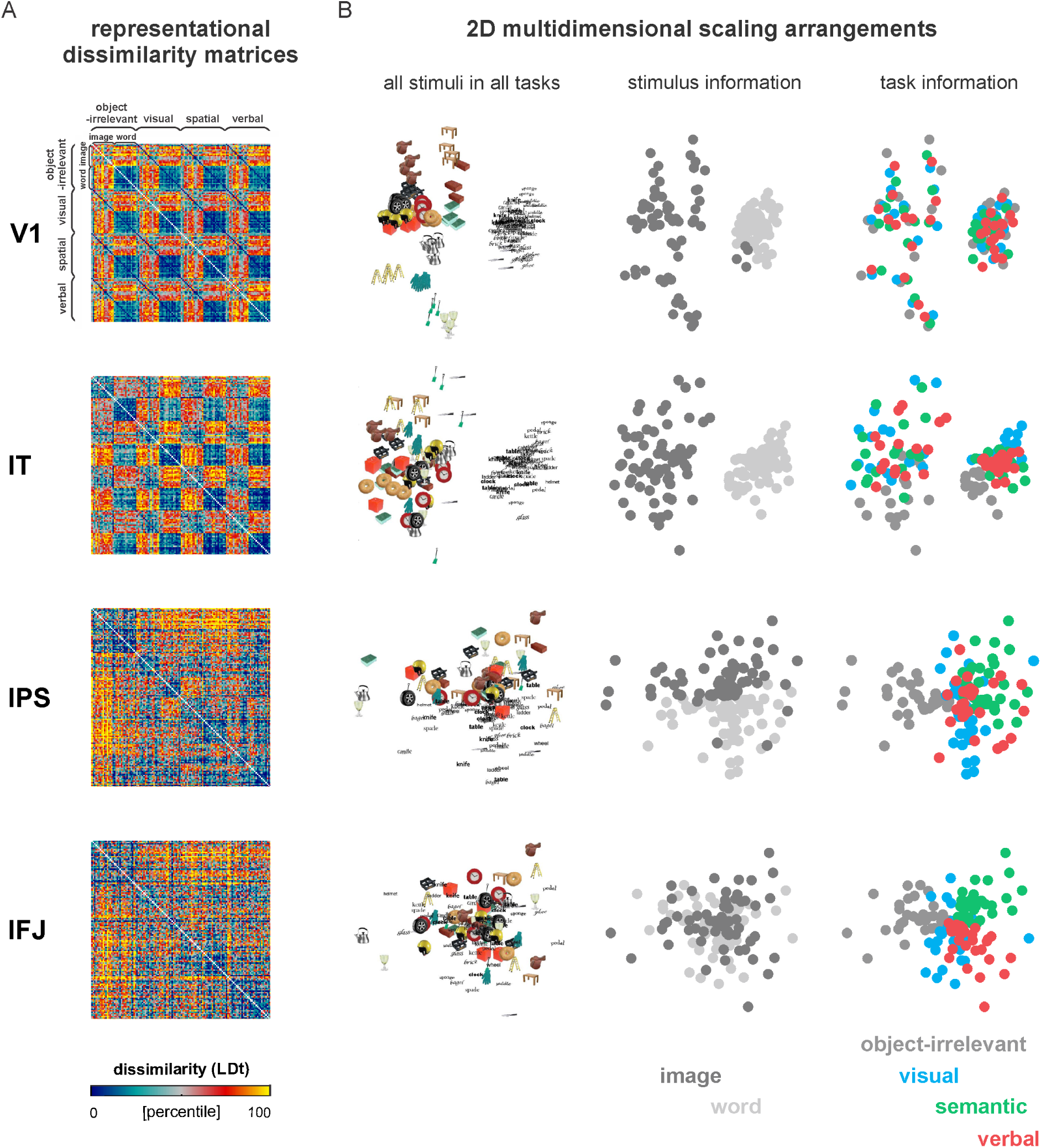
Brain representations of the same stimuli in different task contexts. **A** Representational dissimilarity matrices (RDMs) for the main regions of interest. RDMs contain response-pattern dissimilarities for all condition pairs. Conditions are ordered along the axes by task, and within that, by object. Each RDM is based on data from 14 subjects and two hemispheres, averaged at the level of the dissimilarities (distance measure: linear discriminant t value). The RDMs were independently transformed into percentiles to enable easier comparison between ROIs. IT = inferior temporal cortex, IPS = intraparietal sulcus, IFJ = inferior frontal junction. **B** Multidimensional scaling (MDS; criterion: metric stress) was used to visualize the representation in two dimensions. Distances between stimuli reflect the dissimilarities that are shown in the RDMs: stimuli that elicit similar response patterns are placed close together; stimuli that elicit dissimilar response patterns are placed further apart. From left to right, the MDS arrangements show the actual stimuli in all four tasks, the stimuli intensity-coded by object category, and the stimuli color-coded by task context. The visualisations suggest that the representation in visual cortex (V1, IT) emphasizes stimulus information, showing a clear division between images and words across task contexts. In contrast, the representation in frontal cortex (IFJ) emphasizes task information. The representation in parietal cortex (IPS) appears to emphasize both task and stimulus information.

#### Analysis of representational variance: model selection

We generated a set of repeated-measures ANOVA models to explain the brain data at the level of the RDMs and selected the best model based on performance at predicting held out data. Each model is capable of explaining stimulus, task, and interaction effects. With the current experimental design, there are several configurations for factorizing the conditions (see Figure 3A). The task factor is always fixed at 5 levels: dissimilarities within each of the four tasks: object-irrelevant, visual, semantic, verbal; and dissimilarities between tasks. However, stimulus-related variance can be modeled by a simple one-factor (3 x 1) configuration (dissimilarities within each object category: image, word; and dissimilarities between object categories), or alternatively, more complex two-factor (3 x 3) configurations (image/word/between, distributed across task-relevant object properties: straight/curved/between, indoors/outdoors/between, or one syllable/two syllables/between). If the more complex three-way (5 x 3 x 3) models outperform the simpler two-way (5 x 3) model, this indicates that object properties are represented in a given ROI, possibly more strongly in some tasks than others. If the more complex models do not outperform the simpler model, this indicates that object properties are not represented, or at least not strongly enough to be detected, even when task relevant. We assessed model performance using a leave-one-subject-out cross-validation procedure, fitting the model to all but one subject, and predicting the left-out subject using the parameters of the fitted model. We used built-in matlab functions *fitrm* and *predict*. RDMs were averaged across hemispheres and converted to lower-triangular vectors before model fitting. For each model, performance was estimated by correlating each subject’s predicted RDM with their data RDM using Kendall’s rank correlation coefficient tau a (Nili et al. 2014). We used the subject-average correlation as our test statistic.

**Figure 3.**
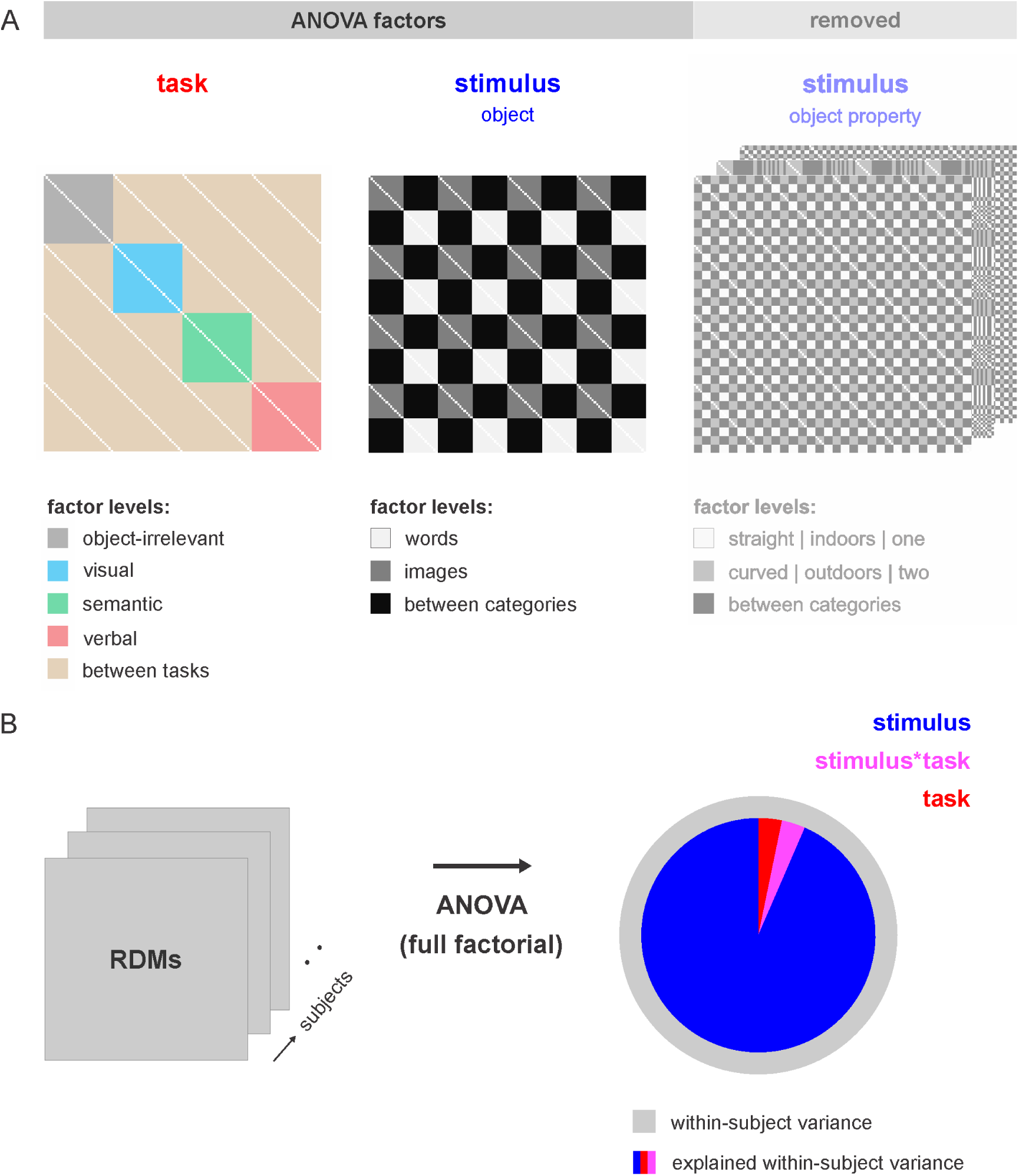
Estimating representational variance components. To determine how strongly task context affects stimulus representations, we partitioned the representational variance into components explained by stimulus, task, and their interaction using a repeated-measures ANOVA across subjects. **A** We first selected the best model from a small set of plausible repeated-measures models, using a leave-one-subject-out cross-validation procedure. Model factors that were considered for inclusion are shown as RDM masks. The masks indicate how dissimilarities were assigned to factor levels. Conditions along the RDM axes are ordered as in Figure 2. A simpler two-way model, which only captures stimulus-related variance at the level of the whole object, performed as well at explaining the brain data as more complex three-way models, which additionally capture stimulus-related variance at the level of object properties (see Figure 3-1). We therefore selected the two-way model for further analysis. **B** We ran a repeated-measures ANOVA for the two-way model using data of all subjects. We estimated the representational variance components for stimulus, task, and their interaction by expressing the sum of squares explained by each of the three model terms as a proportion of the sum of squares explained by the three terms together. The proportions are visualized using a pie chart, showing the relative contribution of each term to the explanatory power of the model (stimulus = blue, task = red, interaction = magenta). The gray circle reflects the total variance associated with the three model terms, both explained and unexplained. The surface area of the pie chart scales with the proportion of explained variance.

We performed statistical inference on the subject-average correlation using a condition-label randomization test (10,000 randomizations), simulating the null hypothesis of no relationship between predicted and data RDMs. If the actual subject-average correlation (for the true labeling) fell within the top 5% of the simulated null distribution (FDR-corrected for multiple models), we rejected the null hypothesis of unrelated RDMs and concluded that model performance was significantly above chance. We then compared performance between each pair of models using bootstrap resampling of the stimuli (1,000 resamplings). This simulates the distribution of differences in subject-average correlation between models that we would expect if we repeated the experiment for different samples of stimuli (drawn from the same hypothetical population of stimuli). If 0 fell in the top or bottom 2.5% of the difference distribution (FDR-corrected for multiple model comparisons), we rejected the null hypothesis of equal model performance. Results are shown in Figure 3-1. Predicted and data RDMs were significantly correlated for all ROIs. However, the three-way models did not outperform the two-way model for any ROI. We therefore selected the more parsimonious two-way model for further analysis.

#### Analysis of representational variance: estimating and comparing variance components

To determine how strongly task context affects the stimulus representations, we partitioned the representational variance into components explained by task (dissimilarities within each of the four tasks: object-irrelevant, visual, semantic, verbal; and dissimilarities between tasks), stimulus (dissimilarities within each object category: image, word; and dissimilarities between object categories) and their interaction using a 5 (task) x 3 (stimulus) repeated-measures ANOVA. Dissimilarities between a stimulus and itself across tasks (Figure 3A, white off-diagonal entries) were not modeled to prevent bias in the stimulus-related variance captured by the model. RDMs were averaged across hemispheres and converted to lower-triangular vectors before model fitting. We fit the model to all subjects and ran a repeated-measures ANOVA for the fitted model using built-in matlab functions *fitrm* and *ranova*. The dissimilarity variance was partitioned using type III sum of squares. Table 4-1 reports inferential ANOVA results, which were corrected for non-sphericity using Greenhouse-Geisser correction.

To enable quantitative comparisons between the variance components estimated by the ANOVA, we developed a measure that indexes the relative contribution of each term to the explanatory power of the model: we express the sum of squares explained by each model term (stimulus, task, interaction) as a proportion of the sum of squares explained by the model terms together. The proportions of explained variance by definition sum to one across model terms. In contrast to the commonly used eta squared, which expresses the sum of squares explained by each model term as a proportion of the total sum of squares, our measure of effect size is independent of model performance. Given that model performance is affected by noise in the data, and that noise levels often differ between ROIs, this is a desirable property that allows descriptive comparisons between ROIs. Similarly, this measure of relative effect size may facilitate comparisons between studies which vary in data quality. We use pie charts to visualize the relative contributions of model terms in explaining representational variance (see Figures 3B and 4). To simultaneously give an impression of model performance, we computed the total sum of squares associated with the three model terms (indexing explained and unexplained variance), and scaled the surface area of the pie chart (indexing explained variance) with respect to this total (see Figure 4). This provides an integrated descriptive visualization of explained and unexplained variance components, given a particular ANOVA model and subject sample. More general indices of model performance can be found in Figure 3-1, where performance is evaluated for predicting individual dissimilarities (as opposed to dissimilarities averaged according to a particular ANOVA model) for subjects left out during fitting.

**Figure 4.**
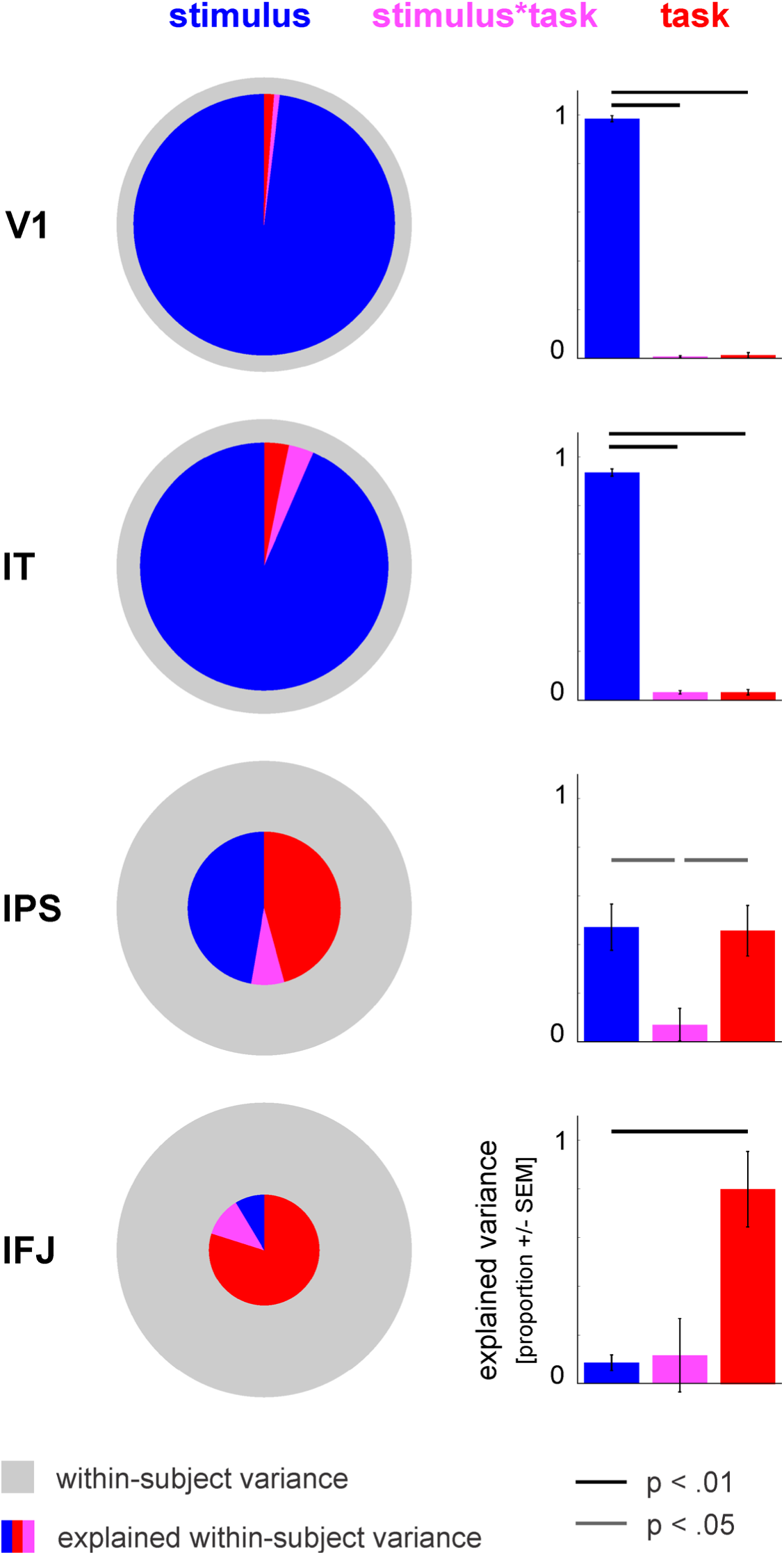
Stimulus effects dominate task effects in visual regions. Pie charts and bar graphs show the relative contributions of stimulus, task, and their interaction in explaining representational variance for the main regions of interest. The explanatory contributions were computed as explained in Figure 3. We tested for differences in the contributions using bootstrap resampling of the subjects (1,000 resamplings, two-sided test). The error bars reflect standard error of the mean computed from the bootstrap resamplings. Significant differences between explanatory contributions are indicated by horizontal bars. Detailed information supporting the displayed results can be found in Figure 4-1 (response-pattern dissimilarities across factor levels) and Tables 4-1 (standard inferential ANOVA results) and 4-2 (relative contributions of stimulus, task, and their interaction for all ROIs). The displayed results indicate that stimulus effects dominate task effects in visual regions, while the reverse is true for prefrontal cortex (IFJ). In parietal cortex (IPS), contributions of stimulus and task are not significantly different. IT = inferior temporal cortex, IPS = intraparietal sulcus, IFJ = inferior frontal junction.

We compared the proportion of explained variance between each pair of model terms using bootstrap resampling of the subjects (1,000 resamplings). This simulates the distribution of differences between proportions that we would expect if we repeated the experiment for different samples of subjects (drawn from the same hypothetical population of subjects). If 0 fell in the top or bottom 2.5% of the difference distribution, we rejected the null hypothesis of equal proportions and concluded that one model term contributes more strongly to explaining the representation than the other. Our approach of estimating and comparing relative variance components can also be incorporated in related analysis frameworks for partitioning multivariate variance (Diedrichsen et al., 2011; Allefeld and Haynes, 2014).

#### Analysis of representational variance: interpretation of effects

Below we describe how main and interaction effects are interpreted using our approach. A main effect of stimulus indicates that dissimilarities between the response patterns elicited by two stimuli depend on the object category of the stimuli. For example, response patterns in the visual ROIs are on average more dissimilar between an image and a word than between two words. A main effect of task indicates that dissimilarities between two stimuli depend on which task they were processed in. For example, response patterns in IPS are on average more dissimilar between tasks than within tasks. This suggests that each task contributes a unique component to the response patterns.

Additionally, task context might strengthen or weaken stimulus information. For example, stimuli might elicit more dissimilar response patterns in one task than in another task. An interaction effect between stimulus and task indicates that the above effects of task depend on the object categories of the stimuli in a pair. For example, in higher-level visual regions V4 and IT, the effects of task are different for image-word pairs than for word-word pairs.

## Results

### Behavioral performance

Subjects performed the tasks on average at 95 percent accuracy. Mean response latency across tasks was 882 ms. We explored potential effects of task and stimulus on accuracy and response latency using a 4 (object-irrelevant, visual, semantic, verbal) x 2 (image, word) repeated-measures ANOVA. For accuracy, we found a significant task x stimulus interaction (F(3,39) = 5.3, p =.0036). Post hoc two-sided paired t-tests revealed different task effects for images than words. For images, accuracy was lower in the object-irrelevant task than in the other three tasks (t(13) =-6.1, p = 3.9 x 10^-5^; t(13) =-4.2, p =.001; t(13) =-6.3, p = 2.7 x 10^-5^; comparison with visual, semantic, and verbal tasks, respectively). For words, accuracy was higher in the verbal task than in the other three tasks (t(13) = 3.6, p =.003; t(13) = 2.7, p =.018; t(13) = 4.2, p =.001; comparison with object-irrelevant, visual, and semantic tasks, respectively). For response latency, we also found a significant task x stimulus interaction (F(3,39) = 44.8, p = 1.0 x 10^-12^). Post hoc two-sided paired t-tests revealed different task effects for images than words. For images, response latencies were longer for the object-irrelevant than the visual task (t(13) = 3.1, p =.0082), and longer for the verbal task than the visual and semantic tasks (t(13) = 5.6, p = 8.8 x 10^-5^; t(13) = 4.9, p = 3.1 x 10^-4^; comparison with visual and semantic tasks, respectively). For words, response latencies were longer for the semantic task than the other three tasks (t(13) = 4.0, p =.0015; t(13) = 3.1, p =.009; t(13) = 4.9, p = 2.7 x 10^-4^; comparison with object-irrelevant, visual, and verbal tasks, respectively).

### Stimulus effects dominate task effects in visual cortex

Figure 2 visualizes the response-pattern dissimilarities for V1, IT, IPS and IFJ. Results for intermediate stages of the ventral visual hierarchy (V2, V3v, and V4) are described below. The RDMs and MDS arrangements in Figure 2 indicate that the representation throughout visual cortex emphasizes stimulus information over task information. Visual ROIs show a clear division between images and words across task contexts. Images appear to elicit more discriminable response patterns than words, consistent with the high degree of visual similarity among words. These observations demonstrate that object category, i.e. whether the stimulus is an image or a word, is a major factor in explaining dissimilarity variance. Importantly, early visual areas V1 and V2 show very similar response patterns to repetitions of the same individual stimulus across task contexts, as indicated by the blue diagonals off the main diagonal. This suggests that early visual representations are largely unaffected by task context. In higher-level visual regions V4 and IT, response patterns are somewhat affected by task context. In particular, the response patterns elicited by the stimuli seen during the object-irrelevant task differ from the response patterns elicited by the same stimuli seen during the other three tasks. A similar effect is observed in V3v. At the same time, however, the representation in V3v also resembles the early visual representation. For example, written words and images of knives, which share low-level visual features, are grouped together. The ‘intermediate’ nature of the V3v representation is consistent with its location along the ventral visual hierarchy. Object properties, i.e. straight/curved, indoors/outdoors, and one/two syllables, do not seem to be strongly represented in visual regions, even when task-relevant, and are therefore not explicitly indicated in Figure 2. In sum, the RDM and MDS visualizations reveal that task effects on stimulus representations in the ventral visual stream are subtle, especially in early visual areas V1 and V2.

To determine exactly how strongly task context affects the stimulus representations, we partitioned the representational variance into components explained by task (dissimilarities within each of the four tasks: object-irrelevant, visual, semantic, verbal; and dissimilarities between tasks), stimulus (dissimilarities within each object category: image, word; and dissimilarities between object categories) and their interaction using a 5 (task) x 3 (stimulus) repeated-measures ANOVA (see Figure 3A). We first ensured that the ANOVA model performed well at predicting response-pattern dissimilarities for held out data (Figure 3-1). Furthermore, all visual ROIs showed significant stimulus effects and most showed significant interaction effects (Table 4-1). We then determined the relative contribution of each model term to the explanatory power of the model, by expressing the variance explained by that term as a proportion of the variance explained by the three terms together. The proportions by definition sum to one across model terms. We therefore use pie charts to visualize the results (see Figures 3B and 4). Stimulus effects account for > 95 percent of explainable representational variance in V1 and V2, and for > 90 percent in V3v, V4, and IT (Table 4-2). For all visual ROIs, the variance component explained by the stimulus term is significantly larger than the variance components explained by the task and interaction terms (bootstrap resampling test; p =.001 for both pairwise comparisons and all visual ROIs; see Figure 4, V2-V4 not shown). In other words, stimulus effects dominate task effects in visual regions.

What drives the stimulus and task effects across the ventral visual stream? As expected from Figure 2, stimulus pairs consisting of an image and a word elicit more dissimilar response patterns than stimulus pairs consisting of two images or two words (two-sided paired t-tests; t(13) > 3.4, p <.0046 for both pairwise comparisons and all visual ROIs). Furthermore, two images elicit more dissimilar response patterns than two words (two-sided paired t-tests; t(13) > 4.6, p < 4.6 x 10^-4^ for all visual ROIs). In other words, response patterns cluster by object category, and clustering is stronger for words than images. All visual ROIs except V3v show significant interaction effects (Table 4-1). The nature of these effects is different for early visual areas V1 and V2 than for higher-level visual regions V4 and IT. In V1 and V2, task mainly affects dissimilarities between images. Images are more discriminable when performing a task at fixation (object-irrelevant task) than when performing a task on the stimulus (visual, semantic or verbal task) (two-sided paired t-tests; t(13) > 2.4, p <.033 for each pairwise comparison). In V4 and IT, task mainly affects dissimilarities between images and words. Images and words are less discriminable when performing a task at fixation than when performing a task on the stimulus (two-sided paired t-tests; t(13) <-2.3, p <.041 for each pairwise comparison). This finding demonstrates that object attention enhances pre-existing category divisions in higher-level visual cortex.

### Task effects dominate stimulus effects in frontal, but not parietal, cortex

In contrast to visual regions, multiple-demand regions IPS and IFJ are visibly affected by task context. The RDMs and MDS arrangements in Figure 2 suggest that the IPS representation emphasizes both stimulus and task information, while the IFJ representation emphasizes task information over stimulus information. We partitioned the representational variance into components explained by task, stimulus, and their interaction using a repeated-measures ANOVA. We first ensured that the ANOVA model performed above chance at predicting response-pattern dissimilarities for held out data (Figure 3-1) and determined that IPS showed significant stimulus and task effects (Table 4-1). IFJ did not show significant effects for individual ANOVA terms (Table 4-1), which is not surprising given the high level of noise in this region (Figure 3-1). However, given that the full ANOVA model performed above chance at predicting held out data (Figure 3-1), we nevertheless compared the relative contributions of the model terms in explaining the response-pattern dissimilarities. For IPS, the relative contributions of task and stimulus do not significantly differ, but both contribute more strongly than the interaction term (bootstrap resampling test; p =.04 for task versus interaction and p =.03 for stimulus versus interaction, Figure 4). For IFJ, task effects account for 80 percent of explainable representational variance. The variance component explained by the task term is significantly larger than the variance component explained by the stimulus term (bootstrap resampling test, p =.002, Figure 4). In other words, task and stimulus effects are similarly strong in IPS, while task effects dominate stimulus effects in IFJ.

What drives the stimulus and task effects in IPS? As expected from Figure 2, stimulus pairs consisting of an image and a word elicit more dissimilar response patterns than stimulus pairs consisting of two images or two words (two-sided paired t-tests; t(13) = 3.4, p =.0048 and t(13) = 6.9, p = 1.0 x 10^-5^, respectively). Furthermore, two images elicit more dissimilar response patterns than two words (two-sided paired t-test; t(13) = 2.8, p =.015). Hence, stimulus effects are similar to those seen in visual cortex, with response patterns clustering by object category, and clustering slightly stronger for words than images. Task effects in IPS are, however, distinct from the task effects observed in visual ROIs. IPS response patterns are more dissimilar between tasks than within each of the four tasks (two-sided paired t-tests; t(13) > 2.9, p <.013 for each within-between task comparison). This suggests that rather than enhancing particular types of stimulus information in IPS, each task context adds a unique component to the response patterns.

## Discussion

We used fMRI and pattern-information analysis in combination with an ANOVA-based modeling strategy to precisely quantify the relative contribution of stimulus and task effects in explaining brain representations throughout the cortical hierarchy. We find that stimulus effects account for the overwhelming majority of explainable representational variance across the ventral visual system: > 95 percent in V1 and V2, and > 90 percent in V3v, V4, and IT. Frontoparietal multiple-demand regions IPS and IFJ exhibit very different results. In IFJ, the relative contributions reverse: task effects dominate stimulus effects, accounting for 80 percent of explainable representational variance. In IPS, contributions of stimulus and task are approximately equal. Our findings demonstrate three main points. First, effects of task on stimulus representations in visual cortex are small, indicating that the visual system favors stability over flexibility. Second, task effects are substantial in multiple-demand regions, suggesting that these regions introduce additional flexibility necessary to support adaptive behaviour. Third, the diverging pattern of results between IPS and IFJ reveals relative specialization within the multiple-demand system.

### From stability to flexibility across the cortical hierarchy

Our findings demonstrate that while task effects are detectable in visual cortex, consistent with prior work (Harel et al., 2014; Kay and Yeatman, 2017; Nastase et al., 2017), the magnitude of these task effects is very small relative to stimulus effects. Several recent studies have indeed reported subtle or undetectable task effects in visual cortex (Bracci et al., 2017; Jackson et al., 2017; Vaziri-Pashkam and Xu, 2017). Our findings suggest that these apparent discrepancies in the literature are attributable to the small effect sizes of task modulations.

The observed minimal contribution of task effects indicates that visual representations are surprisingly stable across task contexts. However, this conclusion seems inconsistent with the dominant view that visual cortex supports flexible, task-dependent representations of stimuli. Is this view solely a consequence of a bias toward publishing positive findings? Not completely. Certain task designs are associated with more clearly observable effects: first, task designs that involve strong competition between stimuli, and thus rely heavily on attentional selection of relevant stimulus information for successful task performance (O’Craven et al., 1999); second, task designs that involve visual search (Peelen et al., 2009; Stokes et al., 2009; Peelen and Kastner, 2011; Çukur et al., 2013). Visual search evokes preparatory response patterns in ventral visual cortex that resemble response patterns elicited by actual targets, such as shapes or people (Stokes et al., 2009; Peelen and Kastner, 2011). These preparatory patterns clearly affect responses to external sensory information (Peelen et al., 2009; Çukur et al., 2013). Hence, tasks that strongly engage attention tend to yield observable modulatory effects in visual cortex. However, while the above studies are suggestive of the dominant view of flexible visual representations, none of them explicitly compare effect sizes of stimulus and task. Notably, they do not provide evidence for complete task dependence of representations. The alternative, consistent with stability of representations, is that stimulus information is reduced when it is task irrelevant, but present nevertheless. Many studies, including several that involve strong competition between stimuli, provide support for the latter possibility (Pinsk et al., 2004; Egner and Hirsch, 2005; Reddy et al., 2009; Bracci et al., 2017; Bugatus et al., 2017). Together, these results indicate that, while task context can modulate stimulus information in visual cortex, representational stability prevails over flexibility.

In contrast to visual regions, parietal and frontal regions show substantial task effects. IPS exhibits an intermediate representation, hosting both stimulus and task information. This finding is consistent with prior work showing that IPS represents both visual information and internal cognitive states (Woolgar et al., 2011; Vaziri-Pashkam and Xu, 2017). IPS especially represents visual information in tasks that are cognitively demanding, involving attention, working memory, or action planning (Silver and Kastner, 2009; Liu et al., 2011; Christophel et al., 2012). Furthermore, studies showing task decoding in IPS typically involve visual stimuli (Zhang et al., 2013; Harel et al., 2014; Vaziri-Pashkam and Xu, 2017). In contrast to IPS, our results indicate that IFJ predominantly hosts task information. Prior work has indeed consistently shown that task context can be decoded from prefrontal cortex (Stiers et al., 2010; Woolgar et al., 2011; Zhang et al., 2013; Harel et al., 2014; Bugatus et al., 2017), in line with its recognised role in cognitive control (Miller and Cohen, 2001; Duncan, 2010). Although we did not detect stimulus information in IFJ, a few previous studies did (Bugatus et al., 2017; Jackson et al., 2017). Importantly, stimulus information in parietal and frontal cortex is often only detectable if it is task relevant (Bracci et al., 2017; Jackson et al., 2017). Together, the IPS and IFJ findings are in line with the multiple-demand hypothesis (Duncan, 2010), which postulates that these regions support flexible cognition and action.

### How does task context affect representations?

Studies typically report that task context selectively affects stimulus information, enhancing information that is task relevant (Peelen et al., 2009; Çukur et al., 2013; Bracci et al., 2017; Nastase et al., 2017). We show a similar result: the image/word category division in higher-level visual cortex is enhanced when fixated objects are task relevant versus when they are not. Our results further suggest that there are limits to the modulatory effects of task context: only stimulus information that is already decodable during a fixation task, namely image information (V1, V2) and image/word category information (V4, IT), is affected. These effects are consistent with the view that primate visual cortex can be modeled as a hierarchical system of relatively stable, increasingly complex image filters (Serre et al., 2007; Güçlü and van Gerven, 2015) that can be selectively strengthened or weakened by higher-order cognitive processes such as attention or working memory. Indeed, cell recordings in nonhuman primates show that attention increases the signal-to-noise ratio of neurons selective for attended image features (Maunsell and Treue, 2006). Furthermore, attention reduces correlated noise between neurons with similar feature preferences (Cohen and Maunsell, 2009; Ruff and Cohen, 2014). In other words, attention not only amplifies but also stabilizes responses to task relevant information in visual cortex (Rabinowitz et al., 2015), biasing population coding toward attended image features among competing visual inputs (Desimone and Duncan, 1995). In higher-level visual cortex, attention tends to select objects as wholes, even when only one object feature is task relevant (Desimone and Duncan, 1995; O’Craven et al., 1999; Vaziri-Pashkam and Xu, 2017).

This is consistent with our finding that the image/word category division in higher-level visual cortex is enhanced if objects are task relevant, even when the tasks focus on other object features.

Task context may not only modulate visual representations through attentional selection of task relevant sensory information. Even in the absence of external stimulation, during tasks involving visual search or working memory, visual population codes can represent task relevant image features (Harrison and Tong, 2009; Peelen and Kastner, 2011). These representations may co-exist or interact with incoming visual information (Rademaker et al., 2018), possibly contributing considerably to previously reported task effects (Peelen et al., 2009; Çukur et al., 2013; Harel et al., 2014; Bracci et al., 2017; Kay and Yeatman, 2017; Nastase et al., 2017; Vaziri-Pashkam and Xu, 2017). Because our study does not include a visual working memory component, object-selective attention is the most likely source of task effects. It is unlikely that differences in response latencies between tasks explain our results because tasks with longer response latencies, and tasks with larger differences in response latencies between images and words, do not yield the largest task effects.

Prior work suggests that modulatory effects of task context on visual representations result from an interplay between top-down signals from parietal and frontal control regions (Gregoriou et al., 2014; Waskom et al., 2014; Kay and Yeatman, 2017), neurochemical signalling (Schmitz and Duncan, 2018), and local canonical computations such as normalization (Reynolds and Heeger, 2009). In our data, these modulatory effects appear to afford only limited representational flexibility, underscoring the importance of higher-level association cortex in introducing further, more profound flexibility.

Neuronal populations in these regions flexibly integrate sensory information with changing behavioral goals (Mante et al., 2013; Rigotti et al., 2013; Fusi et al., 2016). Consistent with a high degree of flexibility, coding of stimulus information in parietal and frontal regions depends heavily on task relevance in both nonhuman primates (Freedman and Assad, 2006; Roy et al., 2010; Mante et al., 2013; Rigotti et al., 2013; Stokes et al., 2013) and humans (Erez and Duncan, 2015; Bracci et al., 2017; Jackson et al., 2017). In the current study, we do not find evidence for task-dependent coding of stimulus information in IPS and IFJ. Task effects are solely driven by coding of task context. This is consistent with the dominance of task coding over stimulus coding in the human fMRI literature on multiple-demand regions (Stiers et al., 2010; Woolgar et al., 2011; Zhang et al., 2013; Harel et al., 2014; Waskom et al., 2014).

## Conclusion

Our study demonstrates that stimulus effects dwarf task effects in visual cortex, while task effects rival or dominate stimulus effects in parietal and frontal multiple-demand (MD) cortex. We conclude that population coding in sensory cortex is optimized for representational stability, providing a consistent interpretation of the external environment for readout by downstream neurons.

Population coding in MD cortex, by contrast, is optimized for behavioral flexibility, representing dynamically changing internal states to provide the necessary basis for intelligent thought and action.

## Acknowledgements

We thank Taylor Schmitz for helpful discussions and comments on the manuscript, and Ian Charest for assistance with defining the IT ROI. This work was funded by the Medical Research Council intramural program (SUAG/002/RG91365 to JD), and by a Dutch Research Council (NWO) Rubicon Postdoctoral Fellowship and a British Academy Postdoctoral Fellowship (PS140117) to MM.

## Data and code availability

The data and code that support the findings of this study will be made publicly available upon peer-reviewed publication.

## Extended data

**Figure 3-1.**
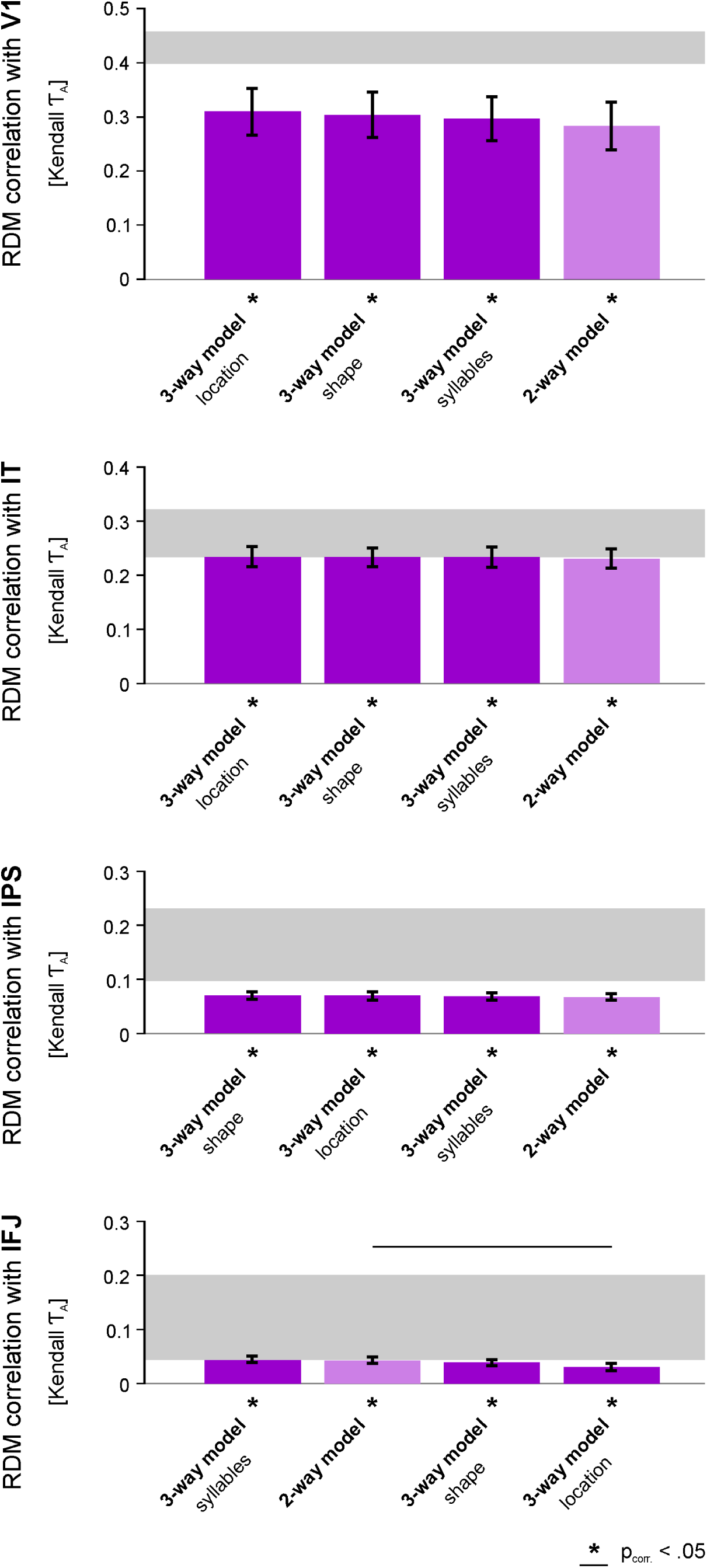
The three-way models do not outperform the two-way model at explaining brain representations. We assessed model performance at explaining brain RDMs for four repeated-measures ANOVA models. The model factors are shown in Figure 3. The two-way model captures variance driven by task context and object category. The three-way models additionally capture variance driven by object-property categories that, depending on the task, are relevant for behavior. We estimated model performance using a leave-one-subject-out cross-validation procedure, fitting the model to all but one subject, and predicting the left-out subject using the parameters of the fitted model. Bar graphs show the correlation between predicted and data RDM, averaged across subjects, for the main regions of interest. The gray bar represents the noise ceiling, which estimates the performance of the true (unknown) model given the noise in the data (Nili et al., 2014). We performed statistical inference on the subject-average correlation using a condition-label randomization test (10,000 randomizations). We then compared performance between each pair of models using bootstrap resampling of the stimuli (1,000 resamplings). The error bars reflect standard error of the mean computed from the bootstrap resamplings. Asterisks indicate significant correlations between predicted and data RDMs. Horizontal bars indicate significant differences in model performance. For each ROI, results were corrected for multiple comparisons by controlling the false discovery rate at.05. The three-way models do not significantly outperform the two-way model for any ROI (V2-V4 not shown). We therefore selected the more parsimonious two-way model for further analysis. IT = inferior temporal cortex, IPS = intraparietal sulcus, IFJ = inferior frontal junction.

**Figure 4-1.**
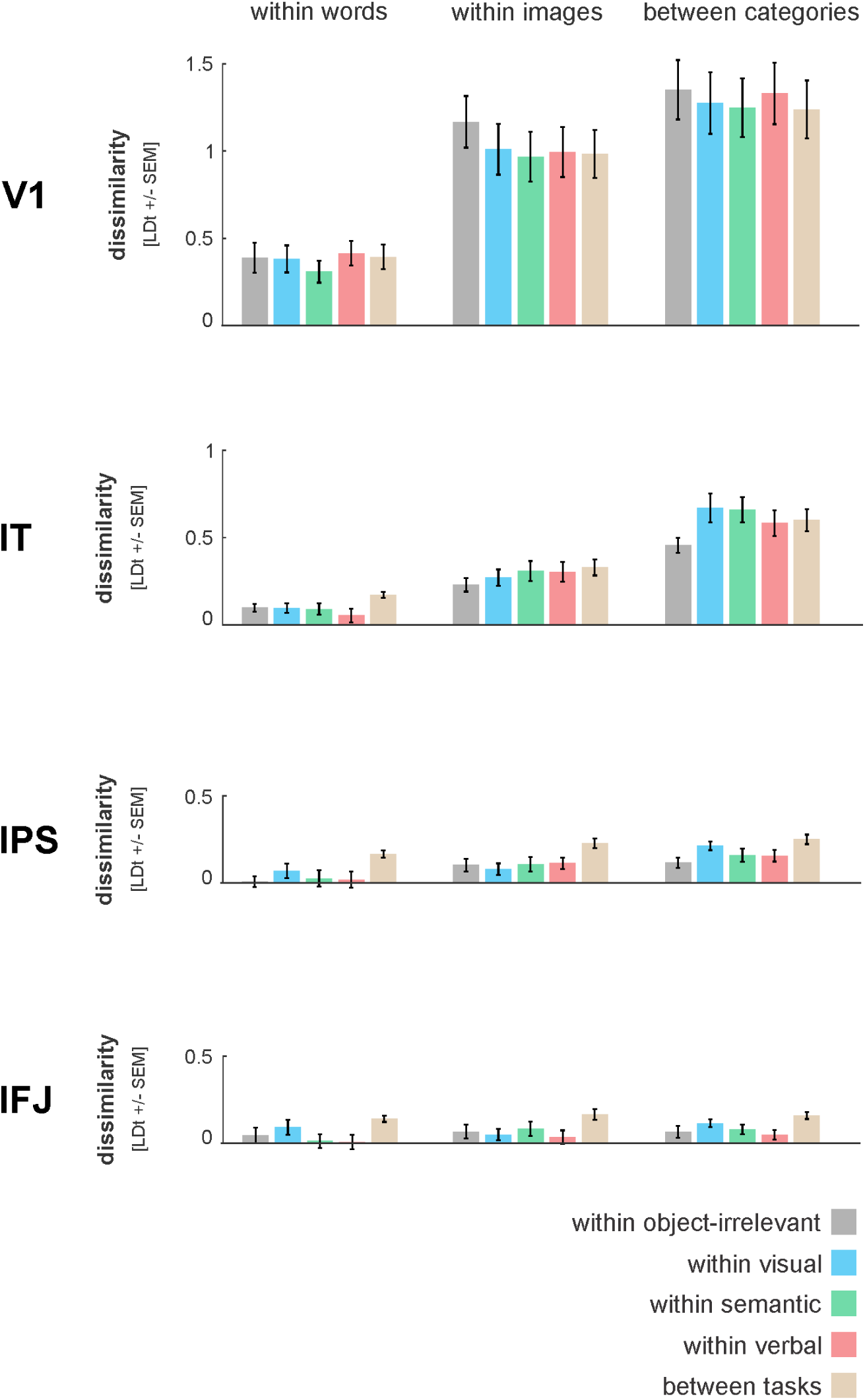
Response-pattern dissimilarities across factor levels for the two-way model. Bar graphs show response-pattern dissimilarities extracted according to the two-way ANOVA model, averaged across subjects, for the main regions of interest. Error bars reflect standard error of the mean across subjects. The single-subject dissimilarities served as input to the ANOVA that yielded the results shown in Figure 4. IT = inferior temporal cortex, IPS = intraparietal sulcus, IFJ = inferior frontal junction.

---

**Table 4-1.** Standard inferential ANOVA results for the two-way model. We used the two-way ANOVA model to partition the dissimilarity variance into components explained by stimulus, task, and their interaction. The dissimilarity variance was partitioned using type III sum of squares. We first performed standard statistical inference on the model terms (results reported in this table) and then determined the relative contribution of each model term to the explanatory power of the model (results displayed in Figure 4 and reported in Table 4-2). The current table lists F values, degrees of freedom, and p values for each ROI and model term. When necessary, p values have been corrected for non-sphericity using Greenhouse-Geisser correction by multiplying the degrees of freedom with the listed epsilon factor Ɛ. Asterisks indicate significant model terms. All ROIs show a significant main effect of stimulus, except for IFJ. IT and IPS also show a significant main effect of task. Furthermore, most visual ROIs show a significant interaction between stimulus and task. IT = inferior temporal cortex, IPS = intraparietal sulcus, IFJ = inferior frontal junction.

|  | stimulus | task | stimulus x task |
| --- | --- | --- | --- |
| **V1** | F(2,26) = 63.5<br>p = $5.7 \times 10^{-7}$ ($\epsilon = 0.57$) ** | F(4,52) = 3.0<br>p = 0.063 ($\epsilon = 0.54$) | F(8,104) = 3.0<br>p = 0.041 ($\epsilon = 0.39$) * |
| **V2** | F(2,26) = 68.0<br>p = $2.4 \times 10^{-7}$ ($\epsilon = 0.59$) ** | F(4,52) = 0.45<br>p = 0.77 | F(8,104) = 4.1<br>p = $9.8 \times 10^{-3}$ ($\epsilon = 0.42$) ** |
| **V3v** | F(2,26) = 43.7<br>p = $4.8 \times 10^{-9}$ ** | F(4,52) = 2.5<br>p = 0.057 | F(8,104) = 1.9<br>p = 0.14 ($\epsilon = 0.45$) |
| **V4** | F(2,26) = 40.4<br>p = $9.6 \times 10^{-7}$ ($\epsilon = 0.71$) ** | F(4,52) = 2.4<br>p = 0.091 ($\epsilon = 0.67$) | F(8,104) = 3.8<br>p = 0.013 ($\epsilon = 0.44$) * |
| **IT** | F(2,26) = 59.7<br>p = $4.7 \times 10^{-7}$ ($\epsilon = 0.60$) ** | F(4,52) = 3.3<br>p = 0.039 ($\epsilon = 0.64$) * | F(8,104) = 5.1<br>p = $3.8 \times 10^{-3}$ ($\epsilon = 0.40$) ** |
| **IPS** | F(2,26) = 19.5<br>p = $6.8 \times 10^{-6}$ ** | F(4,52) = 4.2<br>p = $5.0 \times 10^{-3}$ ** | F(8,104) = 0.99<br>p = 0.41 ($\epsilon = 0.35$) |
| **IFJ** | F(2,26) = 3.4<br>p = 0.085 ($\epsilon = 0.54$) | F(4,52) = 3.0<br>p = 0.074 ($\epsilon = 0.43$) | F(8,104) = 0.63<br>p = 0.58 ($\epsilon = 0.33$) |

\*\* p < .01 \* p < .05

**Table 4-2.** Relative contributions of stimulus, task, and their interaction in explaining representational variance. This table reports proportions and standard error of the mean (SEM) as displayed in Figure 4, for all ROIs. Except rounding errors, proportions sum to 1 across columns. SEM was computed from bootstrap resampling the subjects (1,000 resamplings). IT = inferior temporal cortex, IPS = intraparietal sulcus, IFJ = inferior frontal junction.

|  | **stimulus**<br>proportion [SEM] | **task**<br>proportion [SEM] | **stimulus x task**<br>proportion [SEM] |
| --- | --- | --- | --- |
| **V1** | 0.98 [0.01] | 0.01 [0.01] | 0.01 [0.00] |
| **V2** | 0.99 [0.01] | 0.00 [0.00] | 0.01 [0.00] |
| **V3v** | 0.95 [0.02] | 0.03 [0.02] | 0.02 [0.01] |
| **V4** | 0.94 [0.02] | 0.03 [0.02] | 0.03 [0.01] |
| **IT** | 0.93 [0.01] | 0.03 [0.01] | 0.03 [0.01] |
| **IPS** | 0.47 [0.10] | 0.46 [0.10] | 0.07 [0.07] |
| **IFJ** | 0.09 [0.03] | 0.80 [0.15] | 0.12 [0.15] |

